# Structural and Functional View of Polypharmacology

**DOI:** 10.1101/044289

**Authors:** Aurelio Moya García, Tolulope Adeyelu, Felix A Kruger, Natalie L. Dawson, Jon G. Lees, John P. Overington, Christine Orengo, Juan A.G. Ranea

**Affiliations:** University College London, Institute of Structural and Molecular Biology, London UK; European Molecular Laboratory – European Bioinformatics Institute, Hinxton UK; Department of Molecular Biology and Biochemistry. Universidad de Málaga, 29071 Málaga Spain.; CIBER de Enfermedades Raras (CIBERER), 29071 Málaga Spain.

## Abstract

Protein domains mediate drug-protein interactions and this effect can explain drug polypharmacology. In this study, we associate polypharmacological drugs with CATH functional families, a type of protein domain and we use the network properties of these druggable protein families to analyse their relationships with drug side effects. We found druggable CATH functional families enriched in drug targets, whose relatives are structurally coherent, gather together in the protein functional network occupying central positions, and tend to be free of proteins associated with drug side effects. Our results demonstrate that CATH functional families can be used to identify drug-target interactions, opening a new research direction in target identification.

## Introduction

Systems Pharmacology emerged to address the potential limitations of viewing drug action from the perspective of a single target and has provided some rationale for the need for multi-target approaches in drug discovery^1-3^. The field provides a growing body of evidence against the “magic bullet”, a drug acting on one molecular target, affecting one biological process and thus effecting a cure with few other consequences. Many drugs bind to multiple targets and molecular targets are involved in multiple processes and perform multiple biological functions^4^. Therefore, polypharmacology, the ability of drugs to bind multiple molecular targets and thereby affect multiple biological processes, is likely to be a common phenomenon and can perhaps be harnessed to improve the impact of drug intervention^2,5,6^.

However, polypharmacology is often recognised as an unintended phenomenon^7^, primarily studied from the perspective of side effects, and the rational design of polypharmacological ligands is less frequently undertaken and remains a challenging task^2,8^. Target identification is a crucial task when considering application of polypharmacological compounds and it is important to identify synergistic combinations of targets, rather than single targets^1^. This analysis is often complicated by the fact that many binding events will be silent with respect to phenotypic modulation and emergent drug efficacy.

Most human targets are proteins that are composed of more than one domain^9,10^, but we lack a unified definition of protein domains. In general terms, domains are compact and functional structural units that can be considered the evolutionary and structural building blocks of proteins. Since domains are units of structure11and there is a limited repertoire of domain types^12^, they are combined to form different proteins with different overall functions^13^.

Recent studies have shown that protein domains dominate protein-protein interactions^14^, and mediate the interactions between a drug and its targets^15-17^. It has also been shown that they are a major factor in the polypharmacology of approved and experimental drugs^18^, tend to contain binding sites^17^, and that there are privileged druggable protein domains^19^. These results support the idea that a particular structural domain is likely to be the druggable entity in a protein target. Since proteins have a modular structure and domains recur in different proteins, a reasonable explanation for the fact that a compound binds different protein targets is that they share a domain that is the actual target for the compound.

Under the accepted and general definition that a protein domain is a functional and structural module within a protein, there are several ways to identify and classify protein domains^20^: classification based on structure, SCOP^21^ and CATH^22^; classification based on sequence, Pfam^23^; and function oriented domain classifications such as the functional families classified in CATH, CATH-FunFams^22,24^. CATH-FunFams group together relatives likely to have highly similar structures and functions^25^, and have been highly ranked in the International Critical Assessment of Functional Annotation^26,27^.

In this work, we assess the pharmacology of CATH-FunFams and we explore their ability to direct the biochemical interactions between drugs and their protein targets, since CATH-FunFams group domain targets into families of evolutionary relatives sharing similar structural and functional properties. We found that drug targets are overrepresented in some 81 druggable CATH-FunFams, whose relatives are structurally similar and contain conserved drug binding sites, group together in the protein functional network forming communities, and do not tend to contain proteins associated with drug side effects. Therefore, druggable CATH-FunFams are enriched in potential drug targets rather than off-targets. We propose that CATH-FunFams are a reasonable annotation level for studying the drug-target interactions of polypharmacological drugs, offering valuable insights into possible drug polypharmacology with potential applications in target identification and drug repurposing.

## Results and Discussion

### Drug binding proteins group in druggable CATH-FunFams

Protein domains are classified into protein domain superfamilies when they share a clear evolutionary relationship derived from similarities in their sequence, structure or both. In the CATH classification, a superfamily is sub-classified into functional families (CATH-FunFams) which group domains sharing significant structural and functional similarity. These groupings are achieved by clustering together relatives that have highly similar patterns of sequence conservation and likely specificity determining residues. CATH-FunFams have been benchmarked using experimentally characterised proteins in the Enzyme Classification (EC), SFLD and GO^28^ and have been independently validated by the CAFA independent assessment of function annotation^27^.

There have been a number of efforts to describe the portion of the genome susceptible to interact with drugs conceptualised by the term "druggable genome”, coined by Hopkins and Groom^19^. Despite the use of different definition of protein families (SCOP, Pfam, InterPro) and their different estimates of druggable proteins reported, all the druggable genome studies noted that druggable proteins belong to certain protein families and therefore highlighted the existence of druggable domains^19,29-31^. Therefore, the first step to analyse the role of CATH-FunFams in mediating drug-target interactions was the exploration of the druggable genome they depict.

There are 17229 CATH-FunFams containing 77082 human proteins. Most of them contain only a few protein relatives (the median number of relatives per CATH-FunFam is 3) but a few of them are very highly populated, such as the MHC class I antigen (CATH-FunFam ID 3.30.500.10_3475) which has ca. 14% of the human proteins among its relatives. Using information in ChEMBL^32^, and based on their affinity to bind drugs, we identified a set of 787 human proteins capable of binding drugs (see Methods for details). This set of drug-binding proteins comprise drug targets (i.e. proteins able to bind approved drugs with high affinity) and drug off-targets (i.e. proteins that bind drugs at lower affinities). The drug-binding proteins are distributed in 875 CATH-FunFams (note that many proteins have more than one domain and therefore are represented as relatives of different CATH-FunFams). Most of these functional families are small, containing less than 2% of all human proteins. To have a clear view of the druggable genome captured by the CATH-FunFams, we analysed the proportion of drug-binding proteins across 195 CATH-FunFams that have at least one drug-binding protein among their relatives and contain at least 2% of all the human proteins (Fig. 1). Fig. 1A shows the main druggable protein classes in the 195 CATH-FunFams analysed. Smaller functional families tend to have a higher proportion of drug-binding proteins, although for some druggable classes such as the protein kinases we find very large functional families with a high proportion of drug-binding proteins among their relatives. Fig. 1B suggests that the CATH-FunFams capture well the previously reported druggable genome^19,31^.

**Figure 1.**
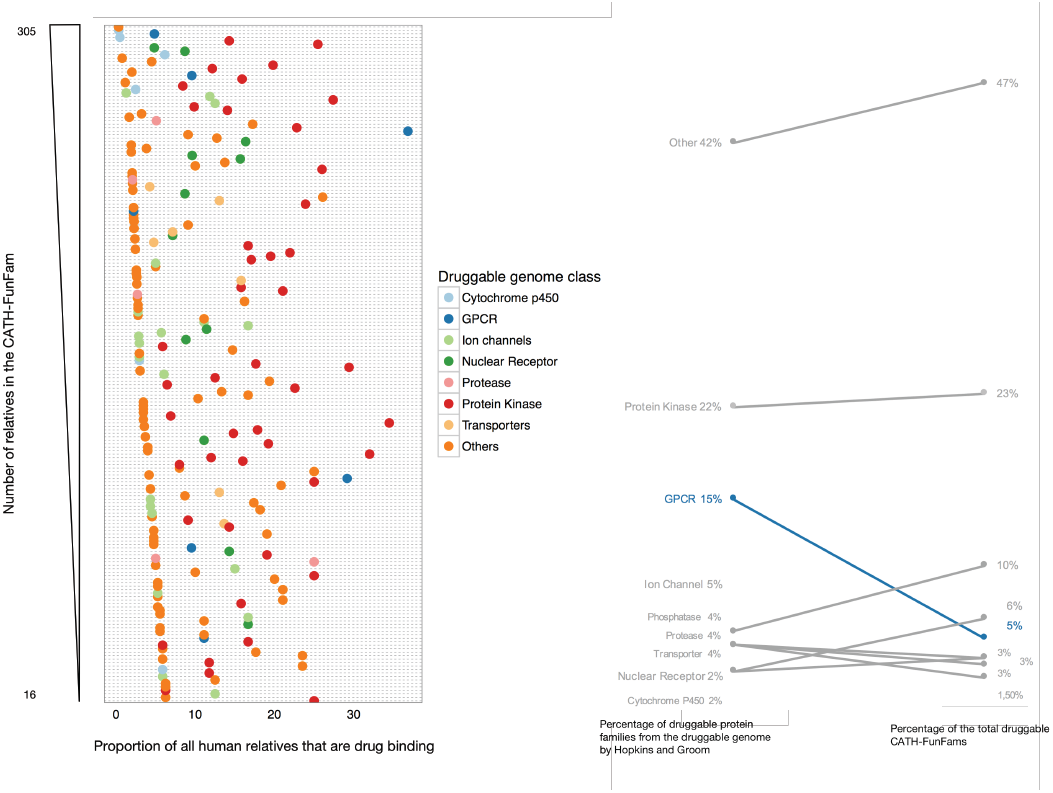
Drug-binding proteins across CATH-FunFams and druggable genome. A) Proportion of drug-binding proteins in CATH-FunFams that have at least one drug-binding protein amongst their relatives and that contain more than 2% of drug targets. B) Slopegraph comparing the previous distribution of druggable protein families (i.e. the druggable genome) by Hopkins and Groom^19^ and our distribution of druggable CATH-FunFams.

Hopkins and Groom defined the druggable genome (i.e. the subset of human proteins able to bind drugs) as a set of 130 InterPro protein families^19^; which were later expanded to an equivalent set of 182 Pfam domains^31^. Although there is no clear equivalence, we see the same main types of druggable domains among the CATH functional families, and the Interpro and PFam families of previous studies: Protein kinases, GPCRs and ion channels cover most of the druggable genome. A recent reassessment of the druggable genome identifies the same privileged druggable proteins families, accounting for 44% of all human targets (GPCRs: 12%; ion channels: 19%; protein kinases: 10%; and nuclear receptors: 3%)^33^. It is interesting to note that the number of GPCRs among the CATH-FunFams analysed is considerably lower than expected based on previous reports of the druggable protein families (see Fig. 1B). One reason for this lies in the difference in purity of functional annotations in CATH-FunFams, compared to annotations in other protein families. CATH-FunFams tend to separate domains according to their functional similarity and multi domain context, and therefore proteins assigned to a single InterPro or Pfam family will be split into several smaller CATH-FunFams. In fact, we find many GPCRs scattered across CATH-FunFams with few relatives, which reflects the diverse functionality of this target category. The lower proportion of GPCRs among the drug-binding CATH-FunFams is also explained by the structural nature of CATH functional families: GPCRs are membrane proteins many of which are structurally uncharacterised and therefore not classified in CATH yet, since CATH requires at least one relative with known structure to initiate a new domain superfamily. This limits the presence of GPCRs in CATH functional families.

### Drug targets are overrepresented in certain CATH-FunFams

As mentioned already above, previous research has shown that drug binding sites are contained within protein domains^17^ and that protein domains mediate drug-target interactions^16^. Furthermore, the recurrent identification of domain families in the druggable genome suggests that the conserved sequence properties and functional similarities within a protein family, are associated with conservation of drug binding sites. This suggests that if one member of the protein family can bind a drug, other members would also be able to bind the same drug or a compound with similar physico-chemical properties^19^. Building on these ideas, we mapped polypharmacological drugs to CATH-FunFams to identify potential new targets and off-targets. In other words, if a drug is associated with a functional family, the relatives of the functional family can be either new drug targets or potential off-targets, leading to undesirable pharmacological effects.

We compiled a drug-target dataset by querying ChEMBL for approved multi-target drugs and the human proteins to which they bind directly at high affinity. For each drug, we computed the statistically significant overrepresentation of their targets for each of the CATH functional families we identified previously in our survey of drug-binding CATH-FunFams (see Methods for details). Our resulting drug to CATH-FunFams mapping gave 359 statistically significant associations (Benjamini-Hochberg false discovery rate p-val < 0.001; see Supplementary Table S1) between 245 approved drugs and 81 CATH-FunFams, for which we will use the term druggable CATH-FunFams. We then investigated our druggable functional families to assess whether they are likely to bind drugs.

### Similar drugs map to the same druggable CATH functional families

We expect that CATH-FunFams mediating the interaction between drugs and targets would share the same characteristics as drug targets. One way to evaluate this is the compliance with the Similarity Property Principle (SPP), which establishes that drugs with similar molecular structure are likely to have the same properties^34^. Since the most relevant drug property is biological activity, produced by interaction with molecular targets^35^, to comply with the SPP a pair of drugs should have similar targets. Therefore, evaluating target similarity for structurally similar drugs is a direct way to compare protein drug targets and CATH-FunFams.

Fig. 2 shows the similarities of the interaction profiles of drugs as a function of their molecular similarity for proteins and CATH-FunFams. For each drug, we determined two different interaction profiles: one is the set of protein targets it binds, and the other is the set of CATH-FunFams that contain the targets of the drug among their relatives. For each drug pair, we compute the similarity of their interaction profile by the Jaccard index. High values of the Jaccard index indicate that a pair of drugs have similar interaction profiles (either protein interaction profiles or CATH-FunFam interaction profiles). Thus, where the association index is 1 the two drugs have the same targets. We observe that for proteins, structurally similar drugs (Tc ≥ 0.65, see Supplementary Fig. S2) tend to have similar interaction profiles (i.e. they tend to bind the same targets) while structurally different drugs bind different targets. Furthermore, when we consider CATH-FunFams this observation is still apparent, suggesting that our drug to CATH-FunFam mapping points to druggable CATH functional families.

**Figure 2.**
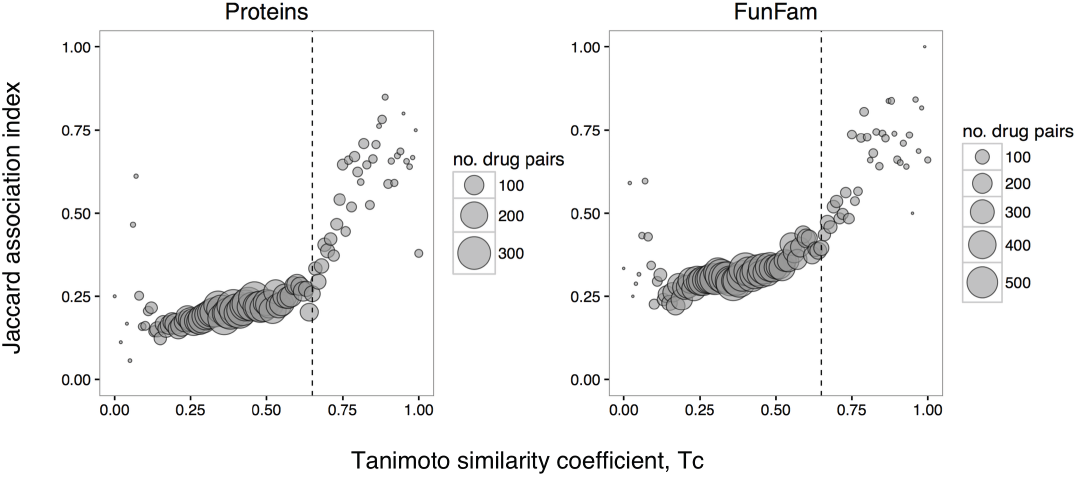
Correlation of the interactions profiles of a drug pair with their molecular similarity. Each circle is the average Jaccard index for the two drug-target datasets at a given bin of Tc similarity (bin size 0.01). The size of the circles is proportional to the number of drug pairs in the corresponding Tc bin. The vertical dashed line indicates the drug similarity threshold, Tc = 0.65 (see Supplementary Fig.S2).

### Relatives in druggable CATH-FunFams are structurally similar with conserved binding sites

Protein targets contain drug-binding sites. Therefore, if our 81 druggable CATH-FunFams are enriched in new potential drug targets, their relatives must contain drug-binding sites. To evaluate the presence of drug binding sites in these 81 druggable CATH-FunFams we examined the 57 CATH-FunFams that have a crystal structure in the PDB, for their enrichment in druggable cavities compared with a set of 100 random non-druggable CATH-FunFams, 63 of them with crystal structure. We found that 75% of the 57 druggable CATH-FunFams with structural information available, have cavities where binding of prodrugs or drug-like molecules is possible. Out of the set of 63 random non-druggable CATH-FunFams with defined structure, only 66% have cavities capable of binding drug-like molecules. Thus, druggable CATH-FunFams have a greater proportion of cavities able to bind drug-like molecules (p-val < 0.0001, Fisher exact test), suggesting that the 81 druggable CATH-FunFams are enriched in potential drug targets.

Since CATH functional families are structurally and functionally coherent, the relatives of a CATH-FunFam that are associated with a drug should have binding sites for that drug. We investigated this by analysing the drugs associated with CATH-FunFams for which there are structures of drug-target complexes in the PDB. Out of the 14 cases we found, we selected 6 examples to illustrate the presence of drug binding sites in the relatives of the druggable CATH-FunFams (Fig. 3). We can see that the drug binding site is very well conserved among the relatives of the functional family, suggesting that all the relatives of a CATH-FunFam that have been associated with a drug, can bind that drug.

**Figure 3.**
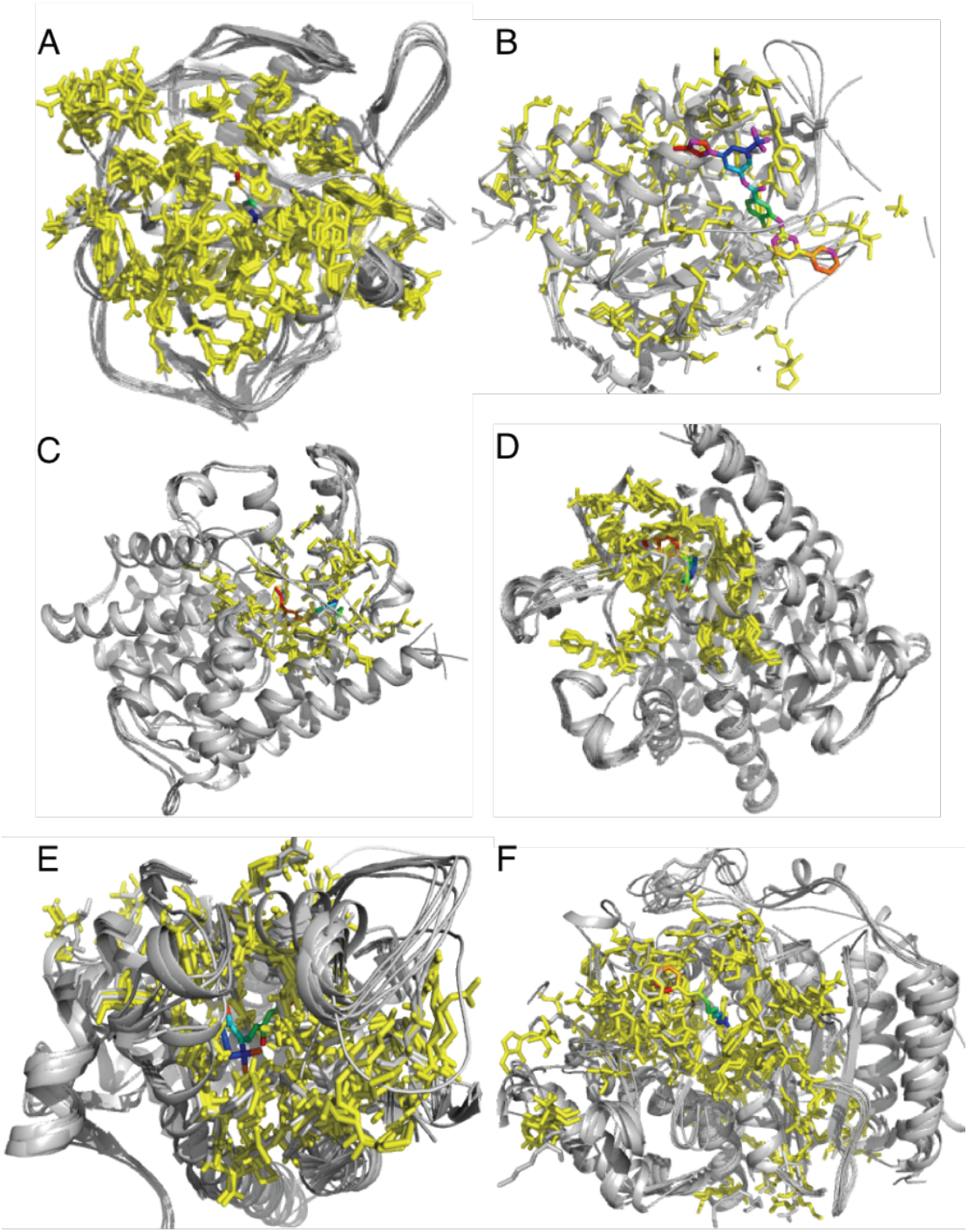
Conservation of the binding site within CATH-FunFams. Structural alignment of the CATH-Funfams associated with: A) acetazolamide (CATH ID: 3.10.200.10-FF1430), B) nilotinib (CATH ID: 1.10.510.10-FF78758), C) Sildenafil (CATH ID: 1.10.510.10-FF78946), D) tadalafil (CATH ID: 1.10.1300.10-FF1260), E) Tretinoin (CATH ID: 1.10.565.10-FF5060) and F) vorinostat (CATH ID: 3.40.800.20-FF2855) and the drug-target complexes of this drugs. The protein domain in all is grey, except the ligand binding residues, which have been mapped across the domains, coloured yellow. The drug molecules are coloured in rainbow.

The mean RMSD for the aligned domain across all the six CATH-Funfams is 1.169 ± 0.812 Å suggesting that the druggable CATH-FunFams are structurally coherent. In order to evaluate the structural conservation of the druggable CATH-FunFams, we clustered the relatives within each druggable CATH-FunFam at 60% sequence identity and aligned the structural representatives of each cluster using the SSAP algorithm^36^. The median RMSD normalised by the number of aligned residues for the 30 druggable CATH-FunFams with structures available is below 5 Å (see Fig. 4), implying that the druggable CATH-FunFams are indeed structurally conserved and that the high conservation of drug binding sites observed in the examples above can be extended to all the druggable CATH-FunFams.

**Figure 4.**
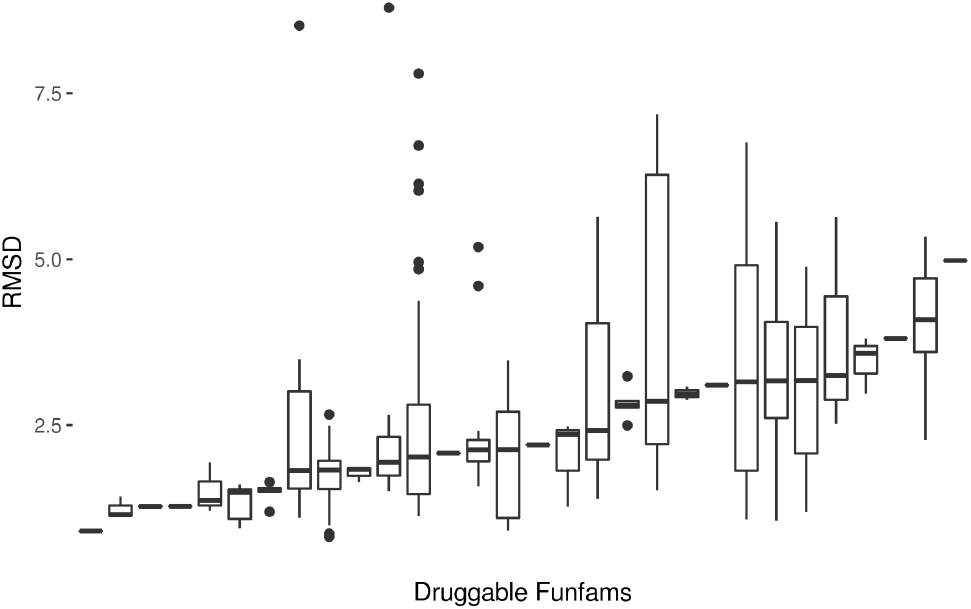
Normalised RMSD of druggable CATH-FunFams. Boxplots of the structural conservation within druggable CATH-FunFams. The RMSD is normalised by the number of residues in each structural alignment.

## Network properties of druggable CATH-FunFams

Molecular systems are modular at different levels, so cellular functions are carried out by modules made up of interacting molecules^37^. Protein functional networks are used to capture this phenomenon – where a link between two proteins means that both are involved in the same function or biological process – and are highly clustered, reflecting this modular design^38^. Therefore, proteins with similar functions tend to be connected or close to each other in the same neighbourhood of the protein functional network^39^. There are many examples of this phenomenon. For example: proteins associated with a disease tend to form modules^40^; modules in protein functional networks are used to predict and uncover protein functions inaccessible to experimental analysis^41,42^; and proteins participating in the same signalling pathway form functional modules^43,44^.

Drug-binding proteins can be either drug targets or off-targets. That is, drugs exert their biological function through interaction with their targets whilst the binding of drugs with off-targets usually results in undesirable side effects^45^. We hypothesised that the targets and the off-targets of a drug may have different network properties and that these may be related in some way to the side effects of a drug.

Since results had indicated that druggable CATH-FunFams contain potential new drug targets and also new off-targets that may lead to undesirable side effects, we analysed the network properties of drug targets and off-targets in order to identify druggable CATH-FunFams that might contain off-targets, leading to undesirable drug side effects.

### Drug targets aggregate in the protein functional network forming neighbourhoods

We analysed the network aggregation of drug targets and off-targets by measuring the mean similarities of the targets of each drug across the kernel transformation of a protein functional network derived from the functional associations between human proteins captured in the STRING database. STRING computes functional associations between proteins through a combined score (ranging from 0 to 1), which indicates the confidence of a given association based on the different types of information supporting that association^46^. We derived a matrix from STRING where the value in row *i*, column *j* had the STRING combined score between protein *i* and protein *j*. This matrix serves the purposes of a similarity kernel^42^, where two proteins with high kernel similarity (i.e. STRING combined score) have a strong connection in the protein functional network.

For each drug, we computed the mean STRING kernel similarity of their targets, off-targets and sets of random proteins with the same number of proteins as the set of drug targets. Figure 5 shows the cumulative distribution function of the STRING kernel similarities for these three datasets. Drug targets have higher STRING kernel similarities than off-targets and both tend to be more similar to each other according to the STRING kernel scores, than expected by chance. This means that drug targets tend to aggregate in the functional network, either forming modules of directly connected proteins or by mapping close to each other in the network, which we dubbed the drug neighbourhood. This tendency of targets to form drug neighbourhoods in the functional network is stronger than off-targets and remarkably larger than expected by chance. We also measured the ability of drug targets to form drug neighbourhoods using the network distance based metrics developed by Menche et al.^40^ and proved, using this alternative approach, that drug targets tend to form drug neighbourhoods regardless of the method used to detect them (see Supplementary Fig. S4).

**Figure 5.**
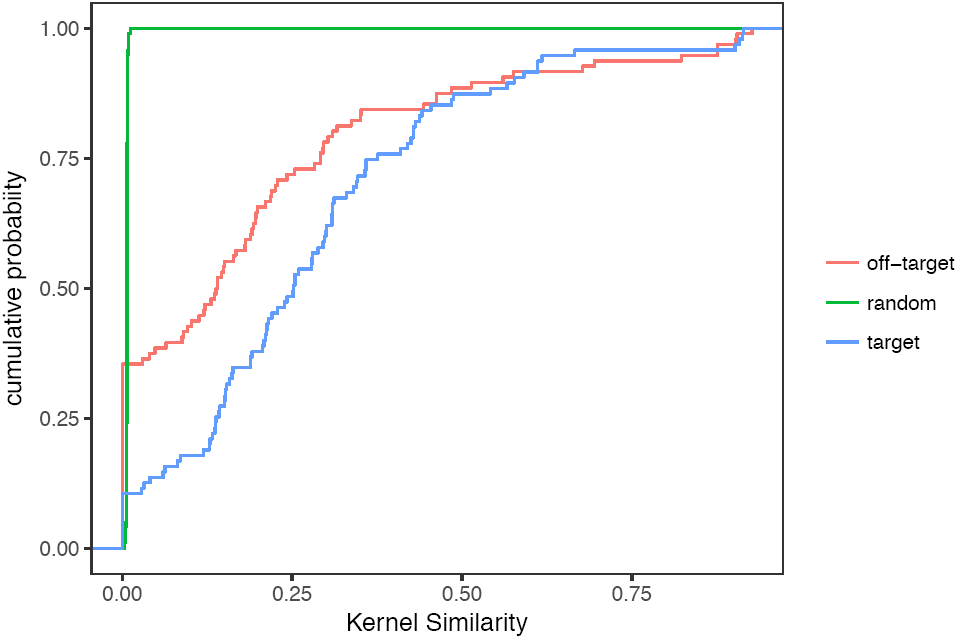
Drug neighbourhoods in the protein functional network. Cumulative distribution function of the kernel similarity of drug targets (blue line), off-targets (red line) and random sets of proteins (green line).

This shows that drug targets tend to cluster in the same neighbourhood of the functional network, whereas off-targets tend to disperse in the network. Since modules in the functional network imply proteins involved in the same process or biological function, we expect that the interaction between a drug and its targets will result in the alteration of one or only a few biological functions (resulting therefore in the drug’s pharmacological effect). By contrast the more dispersed nature of off-targets, which are more likely to be involved in disparate biological processes, will result in many side effects. In other words, proteins binding drugs with many side effects are likely to be more scattered in the functional network whereas proteins binding with less side effects will be more clustered in the functional network.

Our drug-target dataset contains drugs belonging to up to 77 ATC level 2 categories. ATC L01, i.e. antineoplastic agents, is one of the most populated ATC categories in our dataset with 40 drugs. These cancer drugs are an important group that varies widely in the clustering of their targets in the protein functional network (kernel similarities range from 0.00 to 0.93) and their number of side effects (from 13 to 268 adverse drug reactions extracted from SIDER^47^[exp2016-09-19]). Therefore, this is an interesting group of drugs to examine the relationship between drug neighbourhoods in the protein functional network and associated side effects. We observed a strong and significant negative correlation between the kernel similarity of proteins that bind cancer drugs and the number of side effects reported for these drugs in IntSide^48^ (Pearson’s correlation, r = −0.62; p-val < 0.01). The tendency of proteins that bind drugs with many side effects to be dispersed in the functional network holds when we analyse all the drugs in our datasets, although it is weakened by the lack of known side effects data for many of them. This suggests that drug neighbourhoods inform drug safety, and therefore the drug’s potential to affect many biological processes via unintended interactions with other proteins.

### Drug targets are central in the protein functional network

Network centrality is a measure of the importance of certain nodes in the network topology i.e. central nodes are important nodes around which the network revolves. Similarly, for a biological system modelled by a protein functional network, there are essential elements fundamental for maintaining the function of that system. Thus, central nodes of a protein functional network correlate with essential elements of the complex system described by that network^49,50^.

Drug targets have been shown to exhibit a differentiated behaviour on molecular networks, occupying central positions and connecting functional modules^51^. Among the different measures of centrality, betweenness centrality captures best the ability of important nodes to be ‘between’ functional modules and also captures the link between the importance of a node in the network and the essentiality of the protein in the biological system^52^. We used the betweenness centrality measure to assess the importance of drug targets in the protein functional network. Figure 6 shows that drug targets have a higher betweenness centrality than proteins not associated with drugs, represented here by sets of random proteins. However, drug targets are not as central as proteins associated with side effects by IntSide^48^.

**Figure 6.**
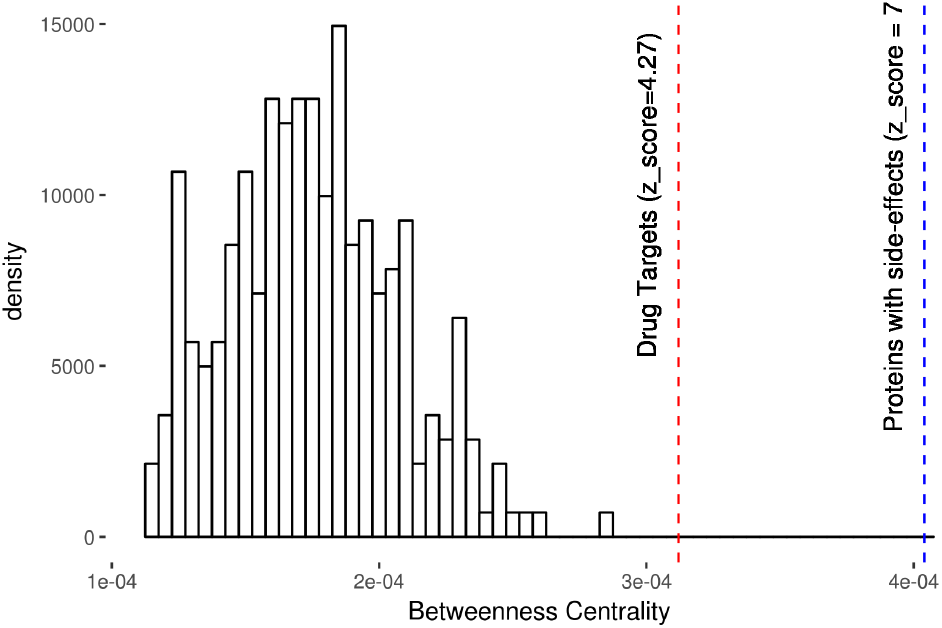
Betweenness centrality of drug targets. The mean betweenness centrality of drug targets (red line) and proteins associated with side effects in IntSide (blue line), in the protein functional network, is compared with the distribution of the mean betweenness centralities of random protein sets.

We observed that drug targets exhibit an interesting nuanced behaviour in the centrality-essentiality continuum: They are important elements in the protein functional network, often bridging two or more modules^51^; however, their essentiality is correlated with the presence of side effects^53^. That is, drug targets occupy central positions in the protein functional network, but if they are highly central (i.e. they are essential) targeting them produces side effects.

### Druggable CATH-FunFams form neighbourhoods in the protein functional network

We extended our analysis to relatives of druggable CATH-FunFams found to mediate drug binding identified by our drug-target association schema. We observed that these are more likely to cluster in the same neighbourhood of the protein functional network (median kernel similarity 0.55) than CATH-FunFams that do not mediate the interactions between drugs and targets (median kernel similarity 0.36). Furthermore, druggable CATH-FunFams can contain relatives that behave as off-targets as well as the relatives that behave as drug-targets. Since the interaction between drugs and off-targets is the main cause of side effects^54^, a CATH-FunFam containing relatives associated with side effects is more likely to include off-targets.

In order to recognise CATH-FunFams, more likely to contain potential targets than off-targets, we built a logistic regression model of the probability of a CATH-FunFam being free of side effect proteins, given its median kernel similarity. According to this statistical model, the probability that a CATH-FunFam which has its relatives completely dispersed on the functional network (i.e. their kernel similarity = 0) does not contain any protein associated with side effects, is 39%. Whilst median kernel similarities greater than 0.43 correspond to an odds ratio greater than 1, that a CATH-FunFams is free of proteins associated with side effects (pval < 0.05). Using this threshold, we can therefore be confident that the relatives of a CATH-FunFam, that gather in the same neighbourhood of the functional network, are more likely to be potential drug targets rather than off-targets and will not be associated with drug side effects.

Furthermore, from our network analyses we previously observed that druggable CATH-FunFams are more likely to be central in the protein functional network than non-druggable CATH functional families (see Fig. 7), and whilst druggable CATH-FunFams have betweenness centralities corresponding to drug targets, they are not enriched in highly central (i.e. essential) proteins whose inhibition would lead to side effects.

**Figure 7.**
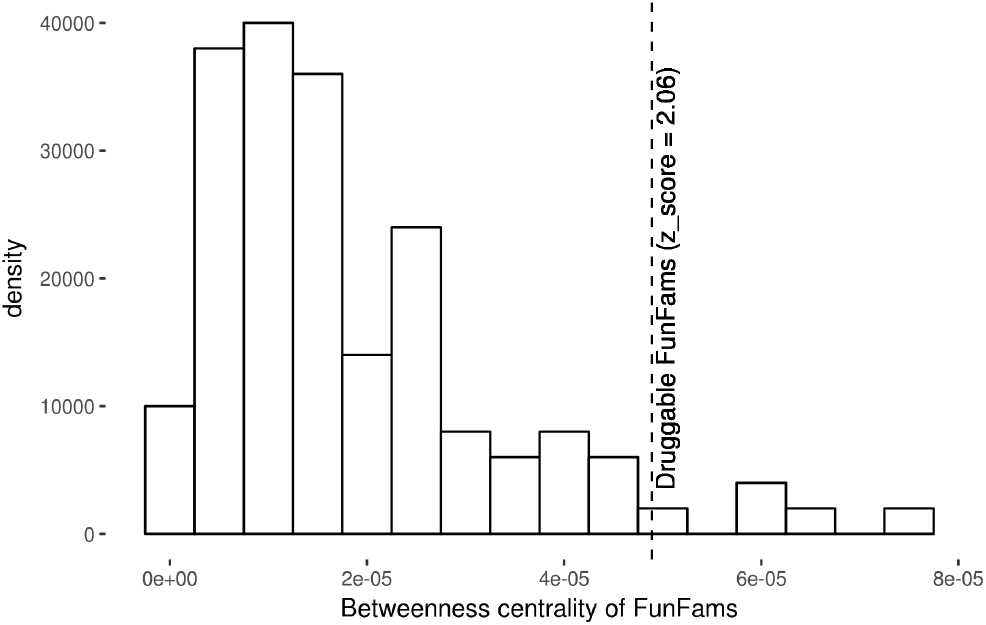
Betweenness centrality of druggable CATH functional families. The mean betweenness centrality of CATH-FunFams (dashed line) is compared with the distribution of the median betweenness centralities of random sets of non-druggable CATH-FunFams in the protein functional network.

### Druggable CATH-FunFams and the polypharmacology and safety of protein kinase inhibitors

Targeted therapies in cancer rely on the inhibition of protein kinases by small-molecule protein kinase inhibitors (PKI) and monoclonal antibodies^33,55^. The superfamily of protein kinases is one of the most important set of drug targets because their dysregulation plays a major causal role in most types of cancer. Kinases are also the largest druggable protein family that bind a common substrate, ATP; so protein kinase inhibitors (PKI) have a great potential for polypharmacology. According to our data, PKI acts on a median number of 28 kinases with high affinity and just 3 of the 37 approved PKI (as of June 2016^33^) are specific to one kinase. Although PKI are considered less toxic than conventional chemotherapeutics, this is not always the case: their broad polypharmacology is a major cause of the observed side effects^55,56^.

We have shown above that druggable CATH-FunFams whose relatives aggregate in network neighbourhoods are likely to be enriched in potential targets and free of off-targets. In fact, we observe a strong correlation between the network spread of CATH-FunFams and the side effects of the PKI associated with them (Pearson’s correlation, r = 0.58; p-val < 0.05). The polypharmacology and mixed safety of PKIs provide an excellent ground to illustrate the use of our druggable CATH-FunFams in spotting potential interactions between drugs and off-targets that lead to side effects. When a kinase inhibitor is associated with a CATH-FunFam that forms a tight network neighbourhood we predict that the inhibitor will have few side effects, conversely a kinase inhibitor associated with a CATH-FunFam much more dispersed in the protein functional network will have many side effects. This is exemplified by the contrast between lapatinib and erlotinib. Lapatinib is a tyrosine kinase inhibitor directed against the oncogenes EGFR and HER2 often used in breast cancer treatment^57^. Lapatinib is regarded as a well-tolerated cancer drug^58^, a characteristic that we could have derived from the association between lapatinib and the CATH-FunFam to which its targets belong and whose relatives form a tightly connected module in the network (see Supplementary Table S1). The median kernel similarity of this CATH-FunFam is 1. In contrast, we associated erlotinib –another EGFR inhibitor used in metastatic non-small cell lung cancer and pancreatic cancer, amongst other types of cancer, and implicated in many severe adverse drug reactions^59^– with a CATH-FunFam whose relatives are more dispersed on the functional protein network (median kernel similarity of 0.22). Therefore, we can anticipate the diversity of side effects caused by erlotinib through the network properties of the druggable CATH functional family associated with it.

Our drug to CATH-FunFams mapping also yields insights into the adverse side effects associated with sunitinib –a receptor tyrosine kinase inhibitor used in the treatment of renal cell carcinoma and other cancers^60^, which has raised safety concerns due to its many adverse reactions^61,62^. We associated sunitinib with two CATH functional families, both very dispersed on the protein functional network (median kernel similarities of 0.16 and 0.15) and hence prone to contain off-targets and cause side effects. Thus, the broad polypharmacology of sunitinib which is associated with functionally diverse targets is captured by our druggable CATH-FunFams. In other words, by targeting more CATH-functional families which are highly spread on the protein functional network, sunatinib causes numerous side effects.

## Conclusion

We have provided fundamental support to the idea suggested by previous research that domain families like CATH functional families provide a useful level of abstraction for a systematic understanding of small molecule bioactivity and drug action^15-18,63,64^. In this work, we conducted further analyses to test whether CATH-FunFams are druggable and have shown that the domains in these families have the potential to be the druggable entities within drug targets. CATH-FunFams comply with the similarity property principle, that is drugs binding domains in these families show a correlation between similarity in their molecular structure and similarity in the targets to which they bind. Furthermore, the functional categories of CATH-FunFams closely agree with the druggable genome reported by other groups examining the functional categories of drug targets. Our studies also examined whether relatives within these functional families tended to be central in the protein functional network (i.e. have high betweenness centrality) and examined whether they were clustered together or highly dispersed in this network. These analyses revealed a greater tendency of druggable CATH-FunFams to be central, than non-druggable CATH-FunFams, and a greater likelihood of relatives in druggable CATH-FunFams to locate close together in the protein network. The value of using CATH-FunFams as proxy targets is further enhanced by the fact that the extent of side effects associated with a drug can be gauged by the dispersion in the protein functional network of relatives in the CATH-FunFams to which the targets of the drug belong.

In summary, our work supports the idea that drug protein interactions are mediated by drug-domain interactions. We have identified the CATH-FunFams as a reasonable annotation level for drug-target interactions, opening a new research direction in target identification with potential applications in drug repurposing.

## Methods

### Drug-proteins dataset

We compiled a drug-protein dataset with 637 drugs and 679 human proteins (including drug targets and off-targets) by querying ChEMBL release 21. ChEMBL allows us to define drug target and drug off-targets based on the concentration at which a drug affects the protein. This provides a way to restrict our dataset to biologically meaningful drug-protein associations. We considered a drug as a small molecule with therapeutic application (THERAPEUTIC_FLAG =1), not currently known to be a pro-drug, reporting a direct binding interaction with single protein (ASSAY_TYPE = ’B’; RELATIONSHIP_TYPE = ’D’; TARGET_TYPE = ’SINGLE PROTEIN’), with a maximum phase of development reached for the compound of 4 (meaning an approved drug). For drug-target interactions we excluded non-specific interactions between small molecules and biological targets by filtering out weak activities (i.e. the activity of a drug against a human protein target should be stronger than 1 μM, where activity includes IC50, EC50, XC50, AC50, Ki, Kd; pchembl_value ≥ 6), while we used a pchembl value threshold between 1 and 4 to capture the less specific interactions between drugs and off-targets^32^.

## CATH-FunFams resource

We used CATH-FunFams v4.1 from CATH-Gene3D v12.0^24,65^. CATH is a protein domain classification system that makes use of a combination of manual and automated structure- and sequence-based procedures to decompose proteins into their constituent domains and then classify these domains into homologous superfamilies (groups of domains that are related by evolution); domain regions in CATH are more clearly defined than in other domain resources by the use of structural data which is more highly conserved than the sequence. CATH-Gene3D is a large collection of CATH^22^ domain predictions for genome sequences ~20 million^66^. CATH superfamilies map to at least 60% of predicted domain sequences in completed genomes using in-house HMM protocols-and as high as 70-80% if more sophisticated threading-based protocols are used^67^. Domain sequences in each superfamily in CATH-Gene3D have been clustered into functionally coherent families (FunFams) using an in-house protocol^28^. This method identifies distinct FunFams within a superfamily having unique patterns of specificity determining residues. CATH-FunFams have been demonstrated to group together relatives likely to have similar structures and functions^28^. They have also been top-ranked in a blind test of functional annotation performance undertaken by the CAFA international assessment^27^.

### Overrepresentation of drug targets in CATH functional families

We evaluated whether the targets 𝕋 {T_1_, … T_*n*_} of a drug *d* are significantly overrepresented among the relatives of a CATH-FunFam ℙ {P_1_, … P_*n*_}. In other words, we want to find whether the CATH-FunFam is enriched in the targets of *d*. For each combination of drug and CATH functional family we defined a test list as the relatives of the CATH-FunFam and a reference list containing all the drug targets. We also defined the expected value for the number of drug targets in the test list, as the number of drug targets that would be expected to be present in the test list based on the reference list. In other words, this is the expected probability that any drug target is a relative of a CATH-FunFam. For example, let’s assume there are a total of 1000 drug targets and 20 of them are relatives of the CATH-FunFam *FF*, then the expected value for *FF* is 0.02 i.e. 2%. If 𝕋 contains 17 proteins we would expect that 0.34 of them are relatives of *FF*, if we observe more than 0.34 targets of *d* among the relatives of *FF*, the targets of *d* are overrepresented on *FF*. Therefore, the overrepresentation of the targets of a drug among the relatives of a CATH-FunFam depends on the expected probability that a protein belongs to the CATH-FunFam. This probability is defined for each CATH-FunFam as the fraction of total drug targets that belongs to the CATH-FunFam.

We calculated a p-value (Benjamini–Hochberg corrected for multiple testing) to determine whether each observed overrepresentation is statistically significant by means of the binomial test. The binomial test evaluates the statistical significance of deviations from the binomial distribution of observations that fall into two categories: (i) the protein is a relative of the CATH-FunFam under consideration, or (ii) the protein is not a relative of the CATH-FunFam under consideration. The binomial distribution is the discrete probability distribution of the number of successes in a sequence of independent yes/no experiments each one with defined success probability. In our case the sequence of independent experiments is 𝕋 the targets of *d*; a success is that a protein from 𝕋 is a relative of the CATH-FunFam under evaluation. Each individual success has a probability 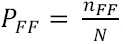, which is the expected probability that a protein is a relative of *FF*, where *n*_*FF*_ is the number of relatives of the CATH-FunFam *FF* and *N* is the total number of proteins relatives of all CATH-FunFams (i.e. all human proteins).

The null hypothesis is that the proteins in 𝕋 are sampled from the same general population as the proteins in ℙ, and thus the probability of observing a target of *d* as a relative of *FF* is the same as observing any protein as a relative of *FF* i.e. *P*_*FF*_. Therefore, the p-value of the binomial test indicates if observing proteins from 𝕋 in the test list ℙ is likely to happen by chance. That is, the p-value for a drug-CATH functional family association indicates the probability of observing the targets of the drug among the relatives of the CATH-FunFam.

For example, let’s consider *FF* with ℙ relatives and *d* with 𝕋 targets, then:

- Success: number of targets from 𝕋 that are in ℙ
- Trials: number of targets of drug *d* in 𝕋
- Probability of success under the null hypotesis: 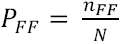
- Overrepresentation threshold: Trials × *P*_*FF*_

The tables below show two examples of overrepresentation of targets of a drug across four CATH-FunFams. For both cases there are 25 possible targets distributed amongst the CATH-FunFams.

1. The drug’s targets belong mainly to one CATH-FunFam

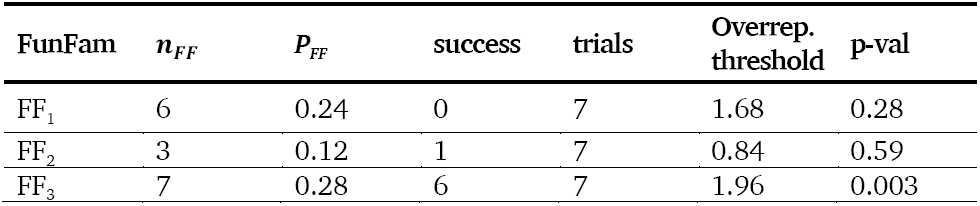

We observe that the targets of the drug are overrepresented for FF_2_ and FF_3_ but the overrepresentation is significant only in FF_3_.

1. The drug’s targets are spread across several FunFams

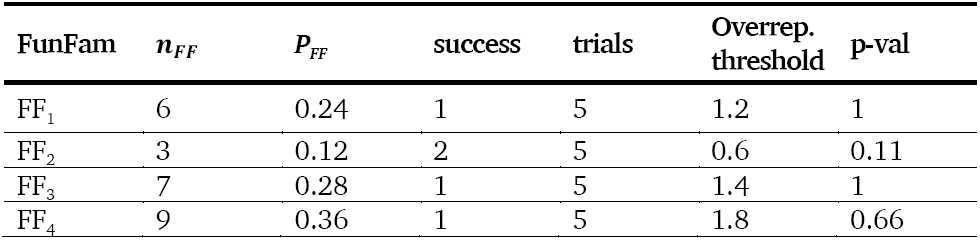

The targets of the drug are overrepresented for FF_2_ but with no statistical significance.

### Molecular similarity calculation and pairwise associations of drug interaction profiles

We performed a significance analysis of the molecular similarity for our set of drugs, in order to choose a threshold Tc which will define a statistically significant level of similarity between any pair of drugs in our dataset. We retrieved the chemical table representing the chemical structure record of 2015 approved drugs (regardless of their targets) from ChEMBL release 21 and we obtained their MACCS molecular fingerprints ^68^. We computed the Tanimoto similarity coefficients (Tc) between each drug and the remaining 2014 drugs using the RDKit package^69^. The Tc similarity quantifies the fraction of features common to the molecular fingerprints of the pair of drugs to the total number of features of the molecular fingerprints of each drug in the pair ^70^. From these distributions of Tc values, we extracted the cumulative distribution function *F*(*t*) that gives the probability of having a similarity less or equal than a given Tc value. A significance level (p-value) defined as *p* = 1 − *F*(*t*) was assigned to every drug for each Tc value, according to Maggiora et al.^34^. Based on this analysis, the Tc threshold to define that two drugs have similar structure is 0.65 (*p* = 0.005; see Supplementary Fig. S2).

For each drug, its interaction profile is the set of targets (proteins or CATH-FunFams) the drug is linked to. We analysed the interaction profile similarity between two drugs by means of the Jaccard association indices (*J*_*ab*_)^71^, defined as 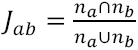 where *n*_*a*_ is the set of elements linked with drug *a* (proteins or CATH-FunFams) and *n*_*b*_ is the corresponding set of elements linked with drug *b*.

### Drug binding sites in CATH-FunFams

We used the Fpocket platform^72^ to detect druggable cavities in the structure of selected domains, i.e. cavities that can bind drug-like molecules. Fpocket is a fast protein pocket prediction algorithm that identifies cavities on the surface of proteins and ranks them according to their ability to bind drug-like small molecules. Thus, Fpocket assesses the ability of a given binding site to host drug-like organic molecules in terms of a druggability scoring function described in ^73^.

To explore whether CATH-FunFams associated with drug binding consist of members with a similar binding pocket and similar amino acid residues, we looked in detail at six examples of FunFams which bind the drugs: acetazolamide, nilotinib, sildenafil, tadalafil, tretinoin and vorinostat. Structural domains from these four different CATH-FunFams were pairwise structurally aligned using SSAP. SSAP scores were used to construct a distance matrix and maximum spanning tree which was then used to derive a multiple superposition of the structural relatives. Data on residues involved in binding each of the drugs of interest were extracted from the NCBI IBIS resource ^74^ using the following PDB IDs as queries: 3ML5 for acetazolamide; 3CS9 for nilotinib; 1UDT for sildenafil; 1UDU for taladafil; 2LBD for treinoin; 4LXZ for vorinostat. These four PDB IDs were chosen as they were the only PDBs in each CATH-FunFam with drug binding information. These drug-binding residue positions were mapped onto the other structural domains using the SSAP alignment data. When producing the figures in PyMOL (www.pymol.org), the number of redundant structural domains in the acetazolamide and vorinostat alignments was reduced to improve clarity.

### Structural coherence of the druggable CATH-FunFams

The structural comparisons of relatives across the druggable CATH-FunFams were done with the SSAP algorithm. Since it is computationally expensive to compare all the relatives of each CATH-FunFam, we analysed the representatives of structural clusters within each CATH-FunFam. Relatives were clustered using CD-HIT^75^ at 60% sequence identity threshold, which indicates significant structural and functional similarity. Representative members of the clusters were used for all-against-all SSAP structural alignments, generating RMSD values normalised by the number of aligned residues in each case.

### Measuring protein neighbourhood in the functional network

We chose STRING to define the protein functional network because it is widely used and frequently updated. STRING compiles protein interaction and functional association data from several sources. These are benchmarked independently, and a combined score (which ranges from 0 to 1) is computed indicating the confidence of the association between two proteins. Therefore, protein associations have higher confidence when more than one type of information supports it^76^, the STRING data can be represented as a network where proteins are linked by their functional and links are weighted by the combined score.

We transformed the STRING network (all edge weights) into a similarity matrix, by taking its adjacency matrix. The adjacency matrix of the full STRING network (i.e. no combined score cut-off) contains all the information of the functional associations between proteins: the value in row *i*, column *j* had the STRING combined score (0-1) between protein *i* and protein *j*. This adjacency matrix has the properties of a Kernel similarity matrix and reflects the integration of the disparate protein interaction types and sources implemented in STRING (see^42,77^ for details on the use of graph-kernels in data integration). Based on this matrix we defined the kernel similarity of a group of proteins as their mean STRING combined score, which reflects the closeness of these proteins in the protein functional network, i.e. proteins with high kernel similarity gather together in the protein functional network.

All data processing, statistics analysis and results plots were produced using Python and Networkx^78^, the R computing environment^79^, and the R library ggplot2^80^.

## Acknowledgements

Aurelio A. Moya-Garcia has received funding from the People Programme (Marie Curie Actions) of the European Union’s Seventh Framework Programme (FP7/2007-2013) under REA Grant Agreement No 623543.

Natalie L. Dawson acknowledges funding from the Wellcome Trust (Award number: 104960/Z/14/Z).

Juan A.G. Ranea was funded by EU-FP7-Systems Microscopy NoE (Grant Agreement 258068), SAF2016-78041-C2-1-R and CTS-486 (Spanish Ministry of Economy and Competitiveness, Andalusian Government and Fondos Europeos de Desarrollo Regional). The CIBERER is an initiative of the Carlos III Health Institute.

The authors thank Dr. Ian Sillitoe and Dr. Tony E. Lewis from the Orengo Group at UCL for their valuable help in obtaining and analysing the structural data; and Dr. Ian Morilla from the Laboratoire Analyse Géométrie et Applications (LAGA), Université Paris 13 - Sorbonne Paris Cité for his help with the statistics analyses.

## Author Contributions

AAMG, CO, and JAGR conceived and designed the study. JPO, JGL and FAK provided constructive suggestions and discussions. AAMG, NLD, JGL and TA performed the experiments and analysed the results. AAMG and CO drafted the manuscript. All authors have read and agreed on the manuscript.

